# Mechanism of NPC1L1 mediated 27-hydroxycholesterol metabolisms in the occurrence and development of osteoporosis

**DOI:** 10.1101/2024.09.30.615783

**Authors:** Bohao Li, Zhicheng Lv, Boyu Chen, Tieqi Zhang, Yueming Yu, Shiwei Sun, Haitian Huang, Lei Zhou, Minghai Wang

## Abstract

Cholesterol metabolism is closely related to the occurrence and development of osteoporosis, but the exact mechanism remains unclear. Niemann-Pick C1-like 1 (NPC1L1) is one of the key cholesterol transporter proteins, however, there are few reports on the functions of NPC1L1 besides regulating cholesterol transport, let alone bone homeostasis. Our preliminary research indicated that NPC1L1 may play a negative regulatory role in osteogenic differentiation. In this study, using in vitro osteogenic differentiation experiment and mouse osteoporosis model, we showed here that NPC1L1 expression was downregulated during osteogenesis, and NPC1L1 knockdown significantly enhanced osteogenic differentiation ability of osteoblasts and delayed progress of osteoporosis. Mechanistically, through RNA sequencing, NPC1L1 was found regulate cholesterol metabolism rather than transportation. It increased 27-Hydroxycholesterol (27-OHC) level through activating 27-hydroxylase (Cyp27a1), resulting in 27-OHC accumulation in osteoblasts and inhibition of osteogenesis. Moreover, C/EBPα was proved mediated NPC1L1 promotes production of 27-OHC by Cyp27a1. These findings reveal that NPC1L1 inhibits osteogenesis and promotes osteoporosis via regulate cholesterol metabolism.

**Graphical Abstract:** NPC1L1 inhibits osteogenic differentiation and promotes the progression of OP through the C/EBPα/Cyp27a1/27-OHC axis.

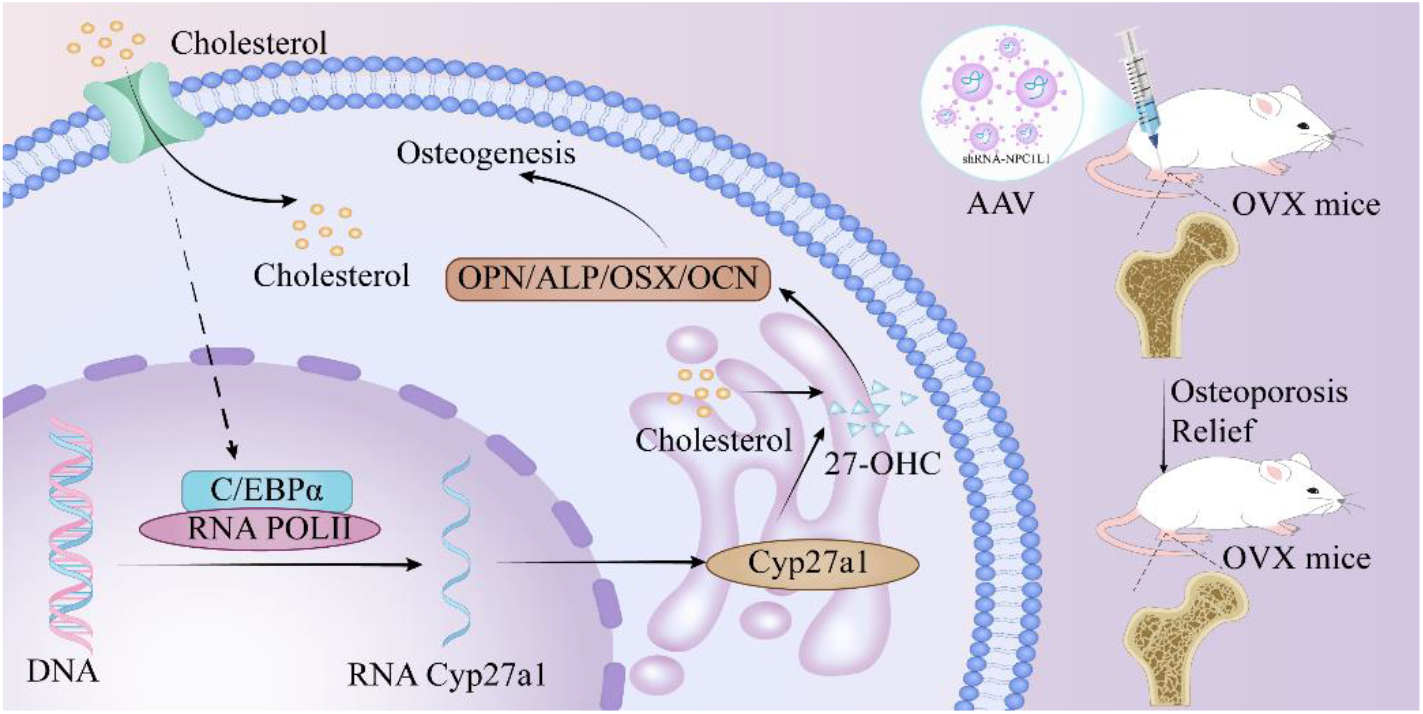

## Introduction

Amidst the global surge in aging populations, the prevalence of osteoporosis (OP) is steadily rising [1, 2]. This condition is marked by diminished bone mass and anomalous microstructure of bone tissue, culminating in heightened skeletal fragility and an elevated susceptibility to fractures. OP represents a systemic metabolic bone disorder [3]. OP development is intricately linked to multiple factors, such as aging, genetic predisposition, unhealthy lifestyle practices, and nutritional deficiencies [4, 5]. Age-related disruption in bone metabolism emerges as a pivotal factor, serving as the primary driver behind the compromised bone formation and heightened bone resorption that collectively underlie the onset of OP [6]. The dynamic integrity of the human skeletal system is maintained through a process known as “bone metabolism”, characterized by bone formation led by osteoblasts and bone resorption dominated by osteoclasts [7]. Due to the intricate nature of bone metabolism mechanisms, the precise molecular intricacies of OP remain elusive. Targeted pharmaceutical interventions are limited, leaving numerous elderly individuals without effective treatment options. Thus, uncovering the exact molecular mechanisms governing bone metabolism is of paramount importance in formulating novel strategies for the prevention and treatment of OP.

Among the diverse differentiation potentials inherent in bone marrow mesenchymal stem cells (BMSCs), the equilibrium between osteogenic and adipogenic differentiation plays a pivotal role in preserving bone homeostasis [8]. Bone homeostasis is intricately linked to bone metabolism [9], the generation of osteoblasts and osteoclasts stands as a pivotal component in the regulation of bone homeostasis [10]. Numerous researchers are actively exploring strategies to enhance the differentiation of BMSCs into osteoblasts or suppress their differentiation into adipocytes. These endeavors seek to offer preventative or therapeutic approaches for OP [11-13]. The prevalence of blood lipid abnormalities often accompanies the aging process in the human body [14]. Moreover, contemporary studies suggest a significant association between lipid pathways and the pathogenesis of OP, Lipid-lowering medications are being explored as potential interventions for the prevention and treatment of OP [15]. The intricate interplay between lipid metabolism and bone metabolism is both captivating and laden with potential.

Cholesterol, a lipid substance, has emerged in recent research as a significant contributor to the occurrence and development of OP as hypercholesterolemia is an independent risk factor for decreased bone mass in postmenopausal women [16]. Cholesterol has been demonstrated to suppress the osteogenic differentiation capacity and proliferation activity of osteoblast precursor MC3T3-E1 cells. Meanwhile, it fosters the formation of osteoclasts and augments their bone resorption functionality [17]. For example, the elevated expression of cholesterol 25-hydroxylase (CH25H), a pivotal enzyme in cholesterol metabolism, in the femurs of ovariectomized mice with osteoporosis provides additional evidence for the regulatory role of cholesterol metabolism in bone metabolism [18]. Furthermore, several clinical studies have consistently demonstrated an inverse association between plasma total cholesterol (TC) levels, LDL-C levels, and bone mineral density (BMD). Notably, when serum HDL-C levels surpass 1.56 mmol/L, there is a notable escalation in the risk of OP incidence [19]. These collective observations indicate that cholesterol metabolism significantly influences the progression of OP. However, the exact molecular mechanisms underpinning this association remain to be fully elucidated.

Niemann-Pick type C1-Like 1 (NPC1L1) is a protein characterized by 13 transmembrane regions, a conserved N-terminal “NPC1 domain,” and a sterol-sensitive domain (SSD). As a critical transporter protein for cellular cholesterol absorption, NPC1L1 is also the target of the cholesterol-lowering drug ezetimibe [20-22]. Moreover, the expression levels of this protein on the cell membrane correlate positively with human blood lipid content [23]. Liu et al. observed that osteopontin (OPN) has the potential to down-regulate the expression of the NPC1L1 gene, resulting in diminished cholesterol absorption and a subsequent reduction in gallstone incidence [24]. However, its specific role in OP warrants further elucidation. In our previous investigation into the gene expression profile of circadian rhythm disrupted BMSCs, we observed a significant association of the NPC1L1 expression with inhibited osteogenic differentiation. In light of these findings, we designed additional cell experiments and observed that the enhanced proliferation and osteogenic differentiation capabilities of NPC1L1 knockdown osteoblasts.

In conclusion, given the existing evidence suggesting a correlation between NPC1L1 expression levels and osteogenic differentiation, we propose that NPC1L1 may play a regulatory role in osteogenesis. To substantiate this hypothesis, our research aims to delineate the precise molecular mechanisms of NPC1L1 modulate bone metabolism, reveal its impact on the occurrence and progression of OP, and assess the therapeutic potential of NPC1L1 in the context of OP.

## Materials and methods

### Materials and Reagents

Alizarin Red S (AR-S, A5533), Alkaline Phosphatase Detection Kit (ALP Stain, SCR004), Dimethyl Sulfoxide (DMSO, D2650), Dexamethasone (DXMS, D4902), L-Ascorbic Acid (AA, A4403), β-Glycerophosphate (β-GP, G9422) and Dodecylpyridinium chloride (CPC, CDS000596) were purchased from Sigma-Aldrich (St. Louis, Missouri, USA). The primary antibodies used included anti-NPC1L1 (NB400-128, Novus, USA), anti-β-actin (AC026, Abclonal, CHN), anti-GAPDH (AC001, Abclonal, CHN), anti-C/EBPα (29388-1-AP, Proteintech, CHN), anti-Cyp27a1 (14739-1-AP, Proteintech, CHN), anti-ALP (A0514, Abclonal, CHN), anti-OPN (22952-1-AP, Proteintech, CHN), anti-OSX (A18699, Abclonal, CHN), anti-OCN (16157-1-AP, Proteintech, CHN), anti-Cyclin d1 (26939-1-AP, Proteintech, CHN), anti-CDK4 (11026-1-AP, Proteintech, CHN), anti-MMP2 (10373-2-AP, Proteintech, CHN), anti-MMP9 (10375-2-AP, Proteintech, CHN). The secondary antibodies utilized were goat anti-rabbit (7074, CST, USA) or anti-mouse (4410, CST, USA) IgG.

### Cell Culture, Osteoblast Differentiation, and Cell Viability Assay via Cell Counting Kit-8 (CCK-8)

The C3H10, C2C12, and 293T (used for lentiviral packaging) cell lines were procured from the Cell Bank of the Chinese Academy of Sciences in Shanghai. C3H10, C2C12, and 293T cells were cultured in high-glucose Dulbecco’s Modified Eagle’s Medium (DMEM, Hyclone) with 10% fetal bovine serum (FBS; Gibco).

For osteoblastic differentiation, the base culture medium was augmented with dexamethasone (DXMS), L-ascorbic acid (AA), and β-glycerophosphate (β-GP) to achieve working concentrations of 0.1 µM, 10 µM, and 10 µM, respectively. Upon reaching approximately 80% confluence, the primary cell cultures were switched to the osteogenic induction medium. All cell lines were incubated at 37 °C in a humidified 5% CO_2_ atmosphere, with medium refreshed bi-daily.

Cell proliferation rate was evaluated using the Cell Counting Kit-8 (CCK-8, Dojindo, Kumamoto, Japan), following the protocol provided by the manufacturer.

### Plasmids and Viral Infection

Plasmids PLKO.1-EGFP-puromycin, PSPAX2, and PMD2.G were acquired from GeneChem (Shanghai, China) to construct a lentiviral packaging system for the expression of short hairpin RNA (shRNA). Three sets of plasmids intended for the suppression of NPC1L1 gene expression were exhibited in Table 1. The full-length coding DNA of NPC1L1 was cloned into the CMV-MCS-EGFP-SV40-Neomycin vector and then transfected into target cells to facilitate overexpression of the NPC1L1.

**Table 1:**
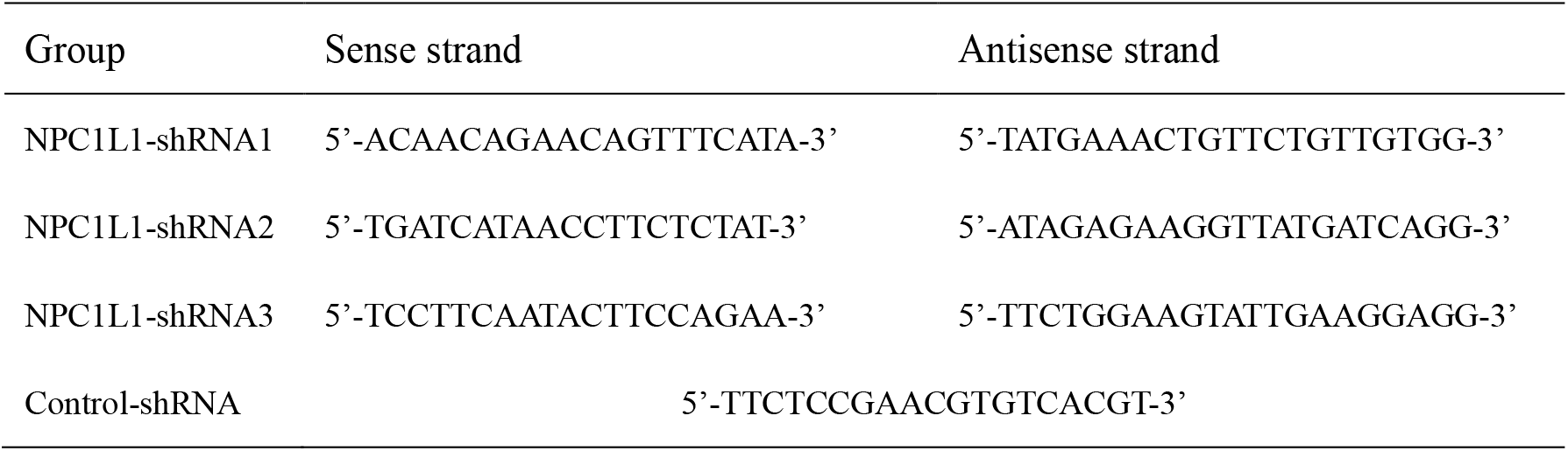
Sequences of Thrap3-shRNA.

For lentivirus production, the components—either pLKO.1-puro or pCDH-CMV-MCS-EF1-Puro vector, psPAX2, and pMD2.G were combined and introduced into 293T cells cultured in high-glucose DMEM devoid of FBS, supplemented with 0.1% LipoFiter (LF-1000, HanBio, Shanghai, CHN). Twelve hours post-transfection, the medium lacking FBS was replaced with fresh medium containing 10% FBS. Supernatants containing lentivirus were harvested at 48 and 72 hours later, filtered through a 0.45 µm cellulose acetate filter (Millipore, Billerica, USA), and aliquots were preserved at -80 °C. Infection of cells was conducted using lentivirus supplemented with 6 µg/mL polybrene. Selection of stably transduced cell lines was achieved using 3 µg/mL puromycin for three days followed by 1 µg/mL for two weeks, thereafter cultivating the cell lines in fresh medium enriched with 10% FBS.

### Alkaline Phosphatase (ALP) and Alizarin Red S (AR-S) Staining

C3H10 and C2C12 cells were seeded in 24-well plates, and osteogenic differentiation was initiated by osteogenic induction medium. On days 0, 3, 6, 9, and 12 of osteogenic induction, culture media were removed and cells were washed twice with phosphate-buffered saline (PBS). Cells were then fixed with 4% paraformaldehyde at 37 °C for 10 minutes. Subsequently, 200 µL of either ALP staining solution (SCR004, Sigma-Aldrich) or AR-S staining solution (A5533, Sigma-Aldrich) was added to each well and incubated at 37 °C for 30 minutes. Post-staining, the wells were rinsed twice with PBS before air drying and documentation through photography. For further analysis of the AR-S staining results, the stained wells were incubated with 100 mM cetylpyridinium chloride (CPC) at 37 °C for 50 minutes. Subsequently, absorbance at 562 nm was measured using a multi-well plate reader (Tecan Infinity 200Pro, Switzerland) to assess the levels of calcification or bone mineral content in the samples.

### Quantitative Real-Time PCR Protocol

Total RNA was extracted employing the RNA-Quick Purification Kit (SB-R001, Share-Bio, CHN), according to the manufacturer’s guidelines. RNA concentration and purity were assessed using an Infinity 200-Pro multi-well plate reader (Tecan, Männedorf, Switzerland). Complementary DNA (cDNA) was synthesized from the total RNA using PrimeScript™ RT Master Mix (RR036A, Takara, JPN) as per the instructions provided.

Quantitative real-time PCR was performed in a 10 µL reaction volume containing 5 µL of 2× SYBR Premix Ex Taq (RR420A, Takara), 1 µL of diluted cDNA, and 0.2 µM of each primer. The amplification protocol involved an initial denaturation at 95 °C for 10 minutes, followed by 40 cycles of denaturation at 95 °C for 15 seconds, annealing at 55 °C for 34 seconds, and a final extension sequence consisting of 15 seconds at 95 °C, 1 minute at 55 °C, 15 seconds at 95 °C, and 15 seconds at 60 °C. Each sample was assayed in triplicate and experiments were independently repeated three times.

Data were analyzed utilizing the 2^−ΔΔCT^ method, with β-actin serving as an endogenous control for normalizing gene expression across samples. Primers for the qRT-PCR were sourced from PrimerBank (https://pga.mgh.harvard.edu/primerbank/), with specifics provided in Table 2.

**Table 2.**
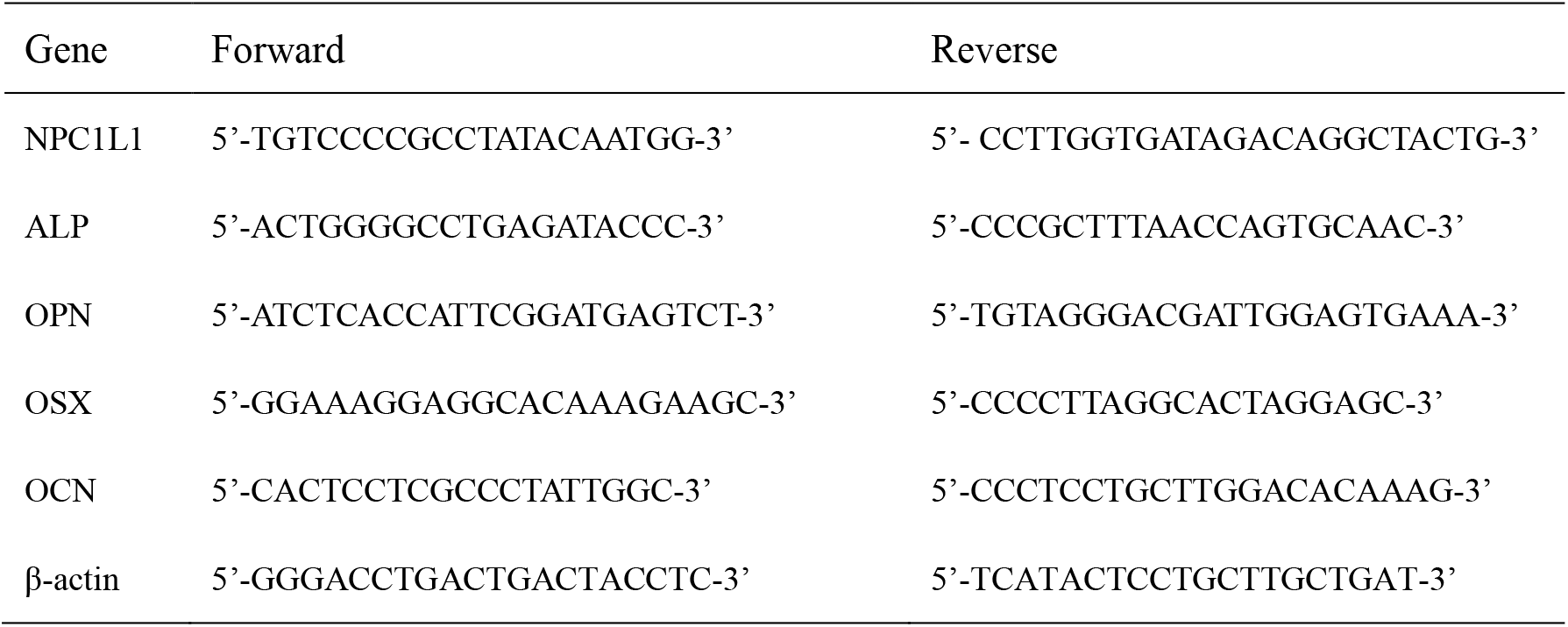
Primer sequences.

### Protein Immunoblotting Procedure

Culture medium was aspirated from 60 mm dishes, and the adherent cells were washed twice with ice-cold PBS. 200 μl of RIPA buffer (p0013b, Beyotime) containing 1% phenylmethylsulfonyl fluoride (PMSF, st506, Beyotime) was added into each dish. Cells were then scraped and the lysates were collected.

Following the instructions provided, the total protein concentration in the supernatant was measured using a BCA Protein Assay Kit (P0010, Beyotime). Subsequently, the protein concentration levels were normalized. Then the protein samples were separated by SDS-PAGE. Following electrophoresis, proteins were transferred onto polyvinylidene fluoride (PVDF) membranes (Millipore, Billerica, MA, USA). Membranes were blocked at room temperature for 20 minutes using Seven Rapid Blocking Buffer (G2052, Servicebio, CHN) and incubated overnight at 4 °C with primary antibodies.

After incubation, membranes were washed thrice with Tris-buffered saline (TBS) containing 0.1% Tween 20 (TBST), each for 10 minutes, and incubated with horseradish peroxidase-conjugated secondary antibodies (7074 and 4410, Cell Signaling Technology, USA) for 2 hours at room temperature. Protein bands were visualized using an enhanced chemiluminescence (ECL) detection kit (Share-bio, China) and imaged with a Fluor Chem E system (ProteinSimple, Santa Clara, CA, USA). Densitometric analysis of the immunoblots was conducted using ImageJ software (version 1.8.0).

### Construction of Animal Models and Group Administration

This animal study was approved by the Animal Welfare and Ethics Group of the Department of Experimental Animal Science at Fudan University (Approval Date: 2023-06-10, Approval Number: 2023-DWYY-24JZS). Specific pathogen-free (SPF) grade female mice, weighing 20-22 g and aged 6 to 8 weeks, were purchased from SPF (Beijing) Biotechnology Co., Ltd. The animals were housed at the Central Experimental Laboratory Animal Facility of the Shanghai Fifth People’s Hospital affiliated with Fudan University.

The mice were randomly divided into four groups: Sham (sham operation group); OVX (ovariectomy group); OVX+Eze (ovariectomy plus Ezetimibe treatment); and OVX+NPC1L1-sh (ovariectomy plus NPC1L1 interfering AAV group). In the OVX, OVX+Eze, and OVX+NPC1L1-sh groups, mice were anesthetized and underwent bilateral ovariectomy along with the removal of surrounding adipose tissue. The Sham group was subjected only to exposure of the ovaries and removal of surrounding adipose tissue before abdominal suturing.

Three weeks post OVX surgery, the mice in the OVX+NPC1L1-sh group received intravenous injections of adenovirus vector fluid to interfere with NPC1L1 expression, while the OVX+Eze group was treated with Ezetimibe and Sham and OVX groups were treated with saline.

### Micro-computed Tomography (Micro-CT) Analysis of Rat Femurs

Following euthanasia, the femurs of the mice were mechanically isolated under aseptic conditions and surrounding soft tissues from the tibia were removed. The bones were then fixed overnight in 4% paraformaldehyde (PFA). The processed femurs were scanned using a high-resolution micro-CT scanner (Bruker), and the images were analyzed using software versions NRvecon 1.6 and CTAn v1.13.8.1. The following six parameters were assessed: Bone volume per tissue volume (BV/TV%), trabecular thickness (Tb.Th), trabecular separation (Tb.Sp), and trabecular number (Tb.N).

### Chromatin Immunoprecipitation

Chromatin immunoprecipitation was performed using the Pierce Chromatin Prep Module (26158, Thermo Fisher Scientific) and Pierce Agarose ChIP Kit (26158, Thermo Fisher Scientific). Cells were cultured to the appropriate density and crosslinked with 1% formaldehyde to fix proteins to chromatin, then quenched with 0.125M glycine. After adding ice-cold PBS containing Halt Cocktail, cells were collected into EP tubes using a cell scraper and centrifuged to remove the supernatant. The pellet was sequentially resuspended in Lysis Buffer 1 and MNase Digestion Buffer Working Solution with protease inhibitors, followed by vortexing, centrifugation, and incubation on ice as needed. Micrococcal Nuclease (ChIP Grade) and MNase Stop Solution were then added sequentially, followed by vortexing, centrifugation, and incubation on ice as needed. The nuclei were subsequently resuspended in Lysis Buffer 2 containing protease/phosphatase inhibitors.After another round of incubation on ice, vortexing, and centrifugation, the supernatant containing the digested chromatin was transferred to a new EP tube. A portion was saved as the input sample. The remaining supernatant was diluted with 1X IP Dilution Buffer, added to a plugged spin column, and combined with the appropriate primary antibody or IgG.After overnight incubation, ChIP Grade Protein A/G Plus Agarose was added, followed by further incubation and centrifugation. The column was retained and sequentially washed with IP Wash Buffers 1, 2, and 3, followed by centrifugation. Agarose beads were resuspended in IP Elution Buffer and incubated at 65 °C with shaking to elute the immunoprecipitated complexes. The eluate was collected. DNA Binding Buffer was added to both the eluate and the saved input sample, and each was transferred to a DNA Clean-Up Column. After centrifugation, the flow-through was discarded.DNA Column Wash Buffer was added, followed by another round of centrifugation and discarding the flow-through. Finally, DNA Column Elution Solution was added directly to the center of each column, and the purified DNA was collected by centrifugation. The resulting solution is the purified DNA. Proceed to PCR or QPCR detection.

### Statistical analysis

SPSS 25.0 (SPSS, Chicago, USA) was used for the statistical analysis. All data were expressed as the mean ± standard deviation (SD). Two groups were compared using Student’s t-test, and more than two groups were compared using one-way analysis of variance (ANOVA). We considered P < 0.05 to be statistically significant.

## Results

### NPC1L1 is down-regulated during osteogenesis and significantly correlated with osteoporosis

In previous research of our team, we verified the circadian clock controls osteoblast differentiation and participates osteoporosis progression, and reveled Cryptochrome 1 (Cry1), one of the core circadian genes, promotes osteogenic differentiation. To further clarify downstream mechanisms, we performed high-throughput sequencing on Cry1 knockdown BMSCs. The gene ontology (GO) analysis of the differentially expressed genes revealed lipid and cholesterol metabolic process were significantly enriched, consistent with previous researches (Figure 1A). Then, the heatmap (Figure 1B) and volcano plot (Figure 1C) of the most significant dysregulated genes showed that cholesterol transporter protein Niemann–Pick C1-like 1 (NPC1L1) was one of the most prominently up-regulated genes, with expression level increased by more than ten times compared to the control group (Figure 1D). Another gene expression profiles of postmenopausal osteoporosis in bone (femur) exhibited that NPC1L1 expression was obviously enhanced, significantly associated with osteoporosis (GSE230665, Figure S1). Subsequently, we conducted preliminary experiment, RT-qPCR verified that NPC1L1 mRNA level was obviously up-regulated in Cry1 silenced osteoblast C3H10 and C2C12 cells (Figure 1E and 1F). Besides, a negative correlation was observed between NPC1L1 expression and osteogenic induction by representative western blot (Figure 1G) and RT-qPCR (Figure 1H) analysis. Besides, there was a significant decrease of NPC1L1 expression after 3 days of induction, indicating an early role in osteogenic differentiation process. Taken together, these data suggest that NPC1L1 is an important inhibitory factor for osteoblast differentiation in osteoporosis.

**Figure 1:**
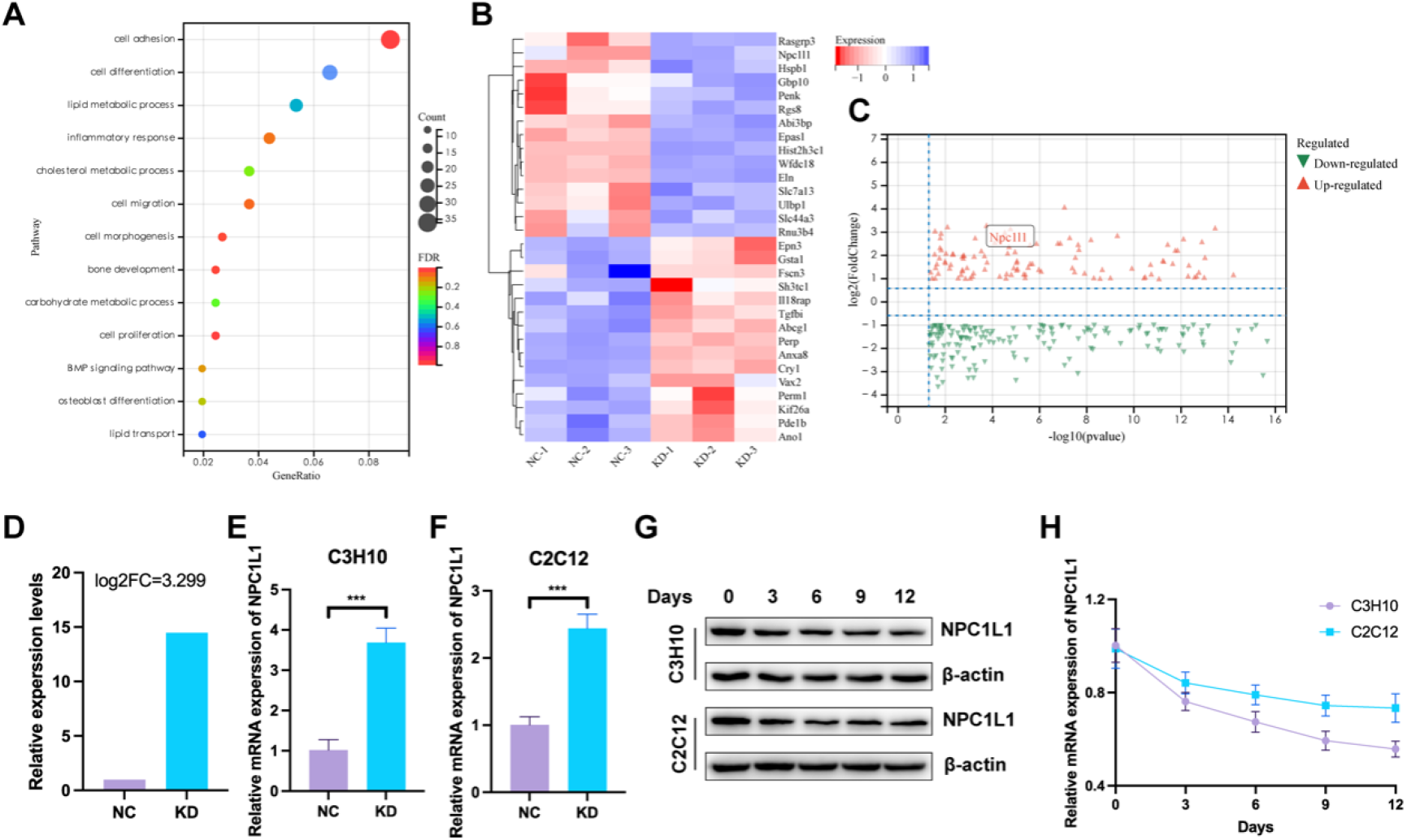
NPC1L1 expression is negatively associated with osteoblast differentiation. A. and B. Gene ontology analysis (A) and heatmap (B) of dysregulated genes identified with RNA-seq in control (NC) and Cry1 knockdown (KD) BMSCs. n = 3 biological replicates. C. and D. Volcano plot (C) and relative expression level (D) of NPC1L1 expression in RNA-seq results. E. and F. RT-qPCR analysis of mRNA expression of NPC1L1 in control and Cry1 knockdown cell lines. G. Representative western blot analysis of NPC1L1 protein level in C3H10 and C2C12 cells treated with osteogenic medium for 0, 3, 6, 9, and 12 days. H. RT-qPCR analysis of mRNA expression of NPC1L1 in C3H10 and C2C12 cells treated with osteogenic medium for 0, 3, 6, 9, and 12 days. All bar graphs are presented as the mean ± SD. *P < 0.05; **P < 0.01; ***P < 0.001.

### Loss of NPC1L1 induces osteogenesis, proliferation, and migration

To determine the exact role of NPC1L1 in osteogenesis and osteoporosis development, lentivirus system was applied to delivery shRNA inserted plasmids into osteoblast C3H10 and MC3T3-E1 cell lines. To ensure the efficiency of NPC1L1 knockdown, three shRNA sequences were designed. Western blotting and RT-qPCR were performed to evaluate the efficiency of these three shRNA in protein and mRNA levels, showing that NPC1L1 was most significantly silenced in NPC1L1-sh group, as both the protein and mRNA levels were most significantly decreased in NPC1L1-sh group in the two cell lines (Figure 2A and 2B). That’s why we chose shRNA-2 for the further analysis (NPC1L1-sh2, written as NPC1L1-sh in the following text).

**Figure 2:**
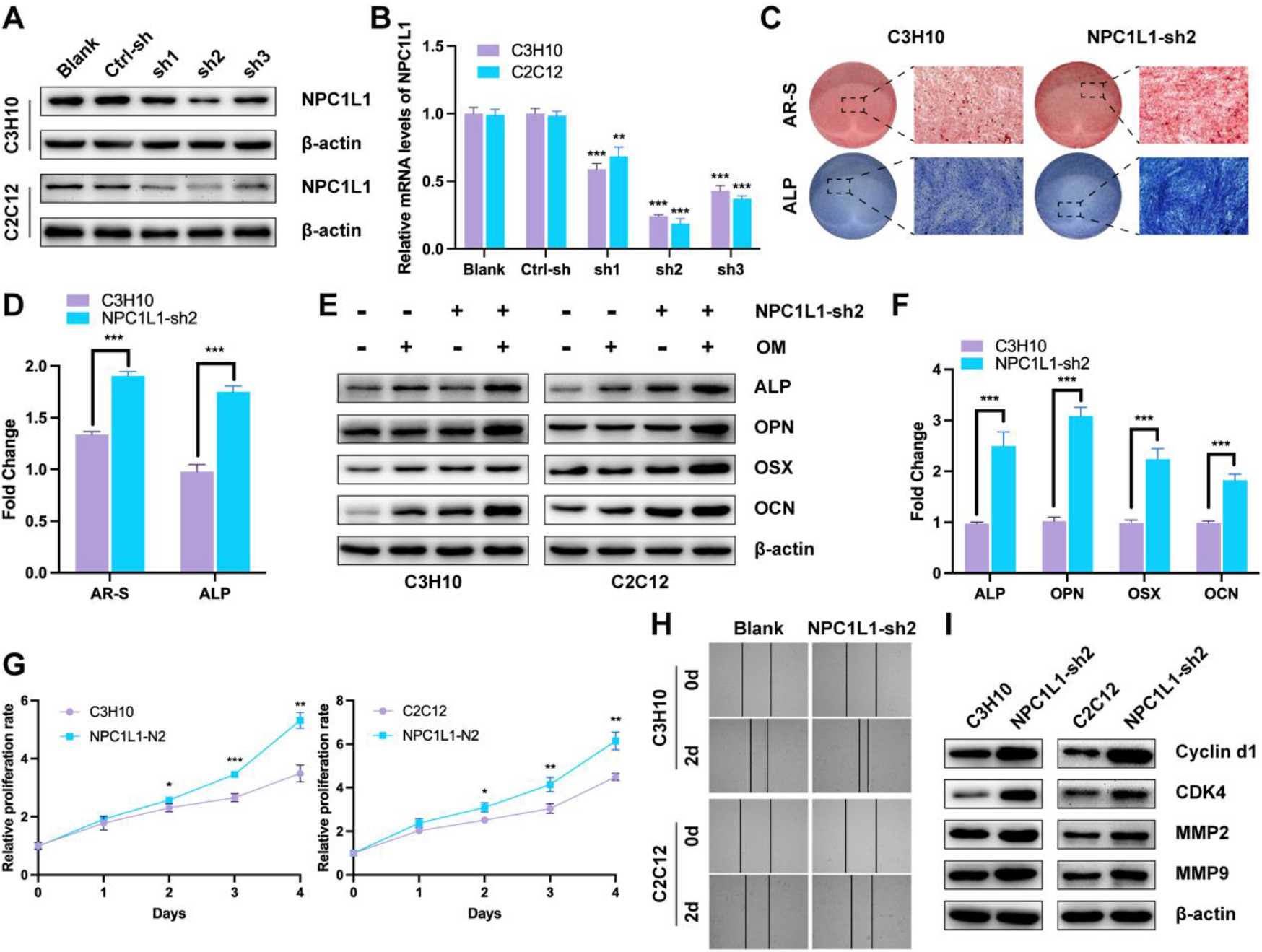
NPC1L1 knockdown enhances osteogenesis, proliferation, and migration of osteoblast cells. A. and B. Representative western blot (A) and RT-qPCR (B) analysis of NPC1L1 knockdown cell lines. C. AR-S (upper panel) and ALP (lower panel) staining of blank and NPC1L1-sh2 C3H10 cells after induction with osteogenic medium for 3 days. D. Quantitative analysis of AR-S and ALP staining results in Figure 2C. E. and F. Representative western blot (E) and RT-qPCR (F) analysis of osteogenic biomarker genes (ALP, OPN, OSX, and OCN) in blank and NPC1L1-sh2 osteoblast cells after induction with osteogenic medium for 3 days. G. and H. CCK-8 (G) and wound healing (H) assays of in blank and NPC1L1-sh2 osteoblast cells. I. Representative western blot analysis of proliferation (Cyclin d1 and CDK4) and migration (MMP2 and MMP9) related biomarkers. All bar graphs are presented as the mean ± SD. *P < 0.05; **P < 0.01; ***P < 0.001.

Next, we observed the role of NPC1L1 in osteoblast differentiation. As expected, osteoblasts response to osteogenic induction, while osteoblasts in NPC1L1-sh group exhibited an increase in Alizarin Red S (AR-S) and ALP staining 3 days after osteogenic induction (Fig. 2C, 2D, and S2). Meanwhile, an elevation of osteogenic biomarkers of ALP, OPN, OSX, OCN protein and mRNA expression were recorded compared to those from the control group (Fig. 2E, 2F), verified that the osteogenic differentiation ability of the NPC1L1-sh group cells was enhanced.

Moreover, during the cell culture process, we found that the growth and migration rates of the NPC1L1-sh group cells were faster than the control group. Therefore, CCK-8 assay and wound healing assay were conducted to test the proliferation and migration rate, respectively. The results showed that the gradient of the cell growth curve in the NPC1L1-sh group was significantly larger, while the scratch healing rate of the NPC1L1-sh group was also significantly higher than that of the control group (Figure 2G and 2H). Accordingly, biomarkers of cell proliferation (Cyclin d1 and CDK4) and migration (MMP2 and MMP9) were increasingly expressed in NPC1L1 knockdown osteoblasts (Figure 2I). These data suggested that NPC1L1 knockdown enhanced the osteogenic differentiation, proliferation, and migration of osteoblasts.

Subsequently, we knocked down NPC1L1 of BMSCs in OVX mice using AAV. 3 months later, the 3D reconstruction images and sectioning of femur by micro-CT showed an increase in the bone formation in the NPC1L1 knockdown mice compared to the WT controls (Figure 3A and 3B). Consistent with this finding, measurement of the BV/TV, Tb.Th, Tb.Sp, and Tb.N revealed a significant increasement in progressive bone formation and delayed osteoporosis (Figure 3C-3F).

**Figure 3:**
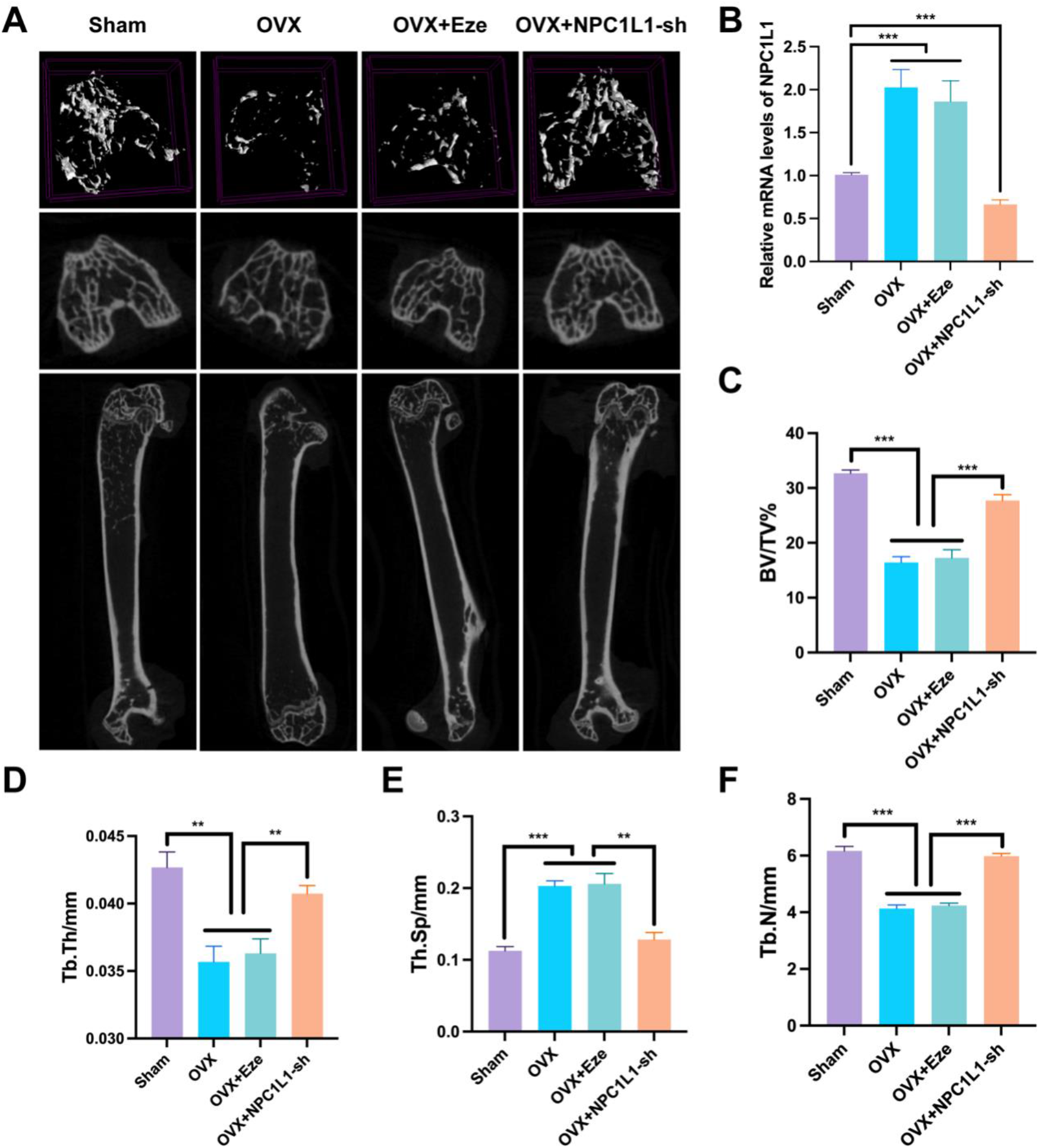
NPC1L1 knockdown delayed the progression of osteoporosis. A. Micro-CT images of femurs of Sham-operated, OVX, OVX+Eze, and OVX+NPC1L1-sh AAV mice. B. Relative mRNA expression in BMSCs from Sham-operated, OVX, OVX+Eze, and OVX+NPC1L1-sh AAV mice. C.-F. Bone volume per tissue volume (BV/TV%), trabecular thickness (Tb.Th), trabecular separation (Tb.Sp), and trabecular number (Tb.N) analysis of Figure 3A. All bar graphs are presented as the mean ± SD. *P < 0.05; **P < 0.01; ***P < 0.001.

### NPC1L1 inhibits osteogenic activity through regulating cholesterol metabolism

We next studied how NPC1L1 controls osteoblast differentiation. We firstly treated C3H10 and C2C12 cells with ezetimibe (Eze), the NPC1L1 specific inhibitor of cholesterol transport. The AR-S and ALP staining (Figure 4A and 4B), as well as the mRNA expression of osteogenic biomarkers ALP, OPN, OSX, and OCN (Figure 4C), showed there was no significant differences between the two groups after induction, indicating Eze has no effect on the cells’ osteogenic ability.

**Figure 4:**
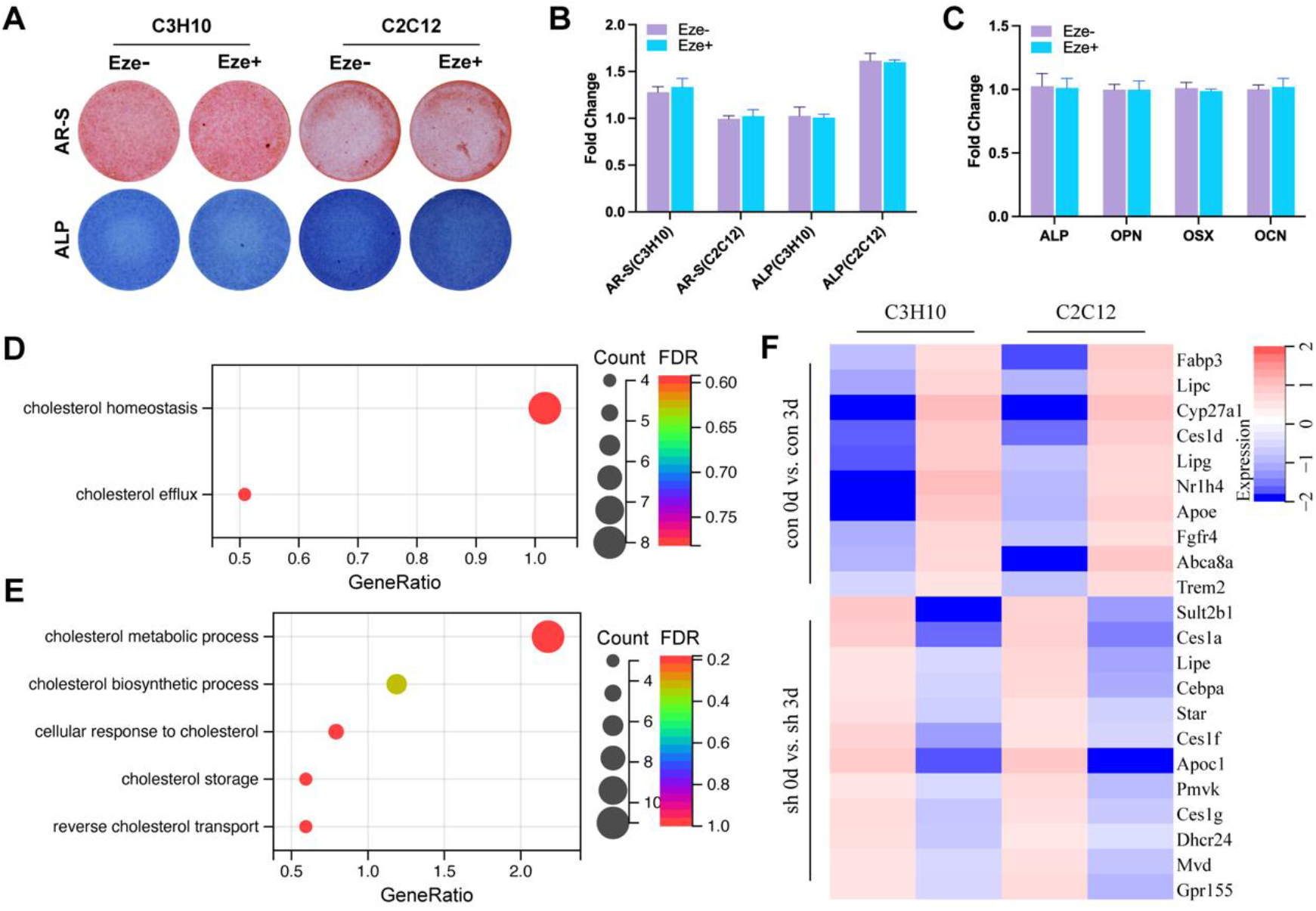
NPC1L1 regulates osteoblast differentiation through cholesterol metabolism. A. AR-S (upper panel) and ALP (lower panel) staining of C3H10 and C2C12 cells treated with or without NPC1L1 inhibitor Ezetimibe. B. Quantitative analysis of AR-S and ALP staining results in Figure 3A. C. RT-qPCR analysis of osteogenic biomarker genes (ALP, OPN, OSX, and OCN) in C3H10 cells treated with or without NPC1L1 inhibitor Ezetimibe. D. Gene ontology analysis of up-regulated genes identified with RNA-seq in blank C3H10 cells after induction with osteogenic medium for 3 days. E. Gene ontology analysis of down-regulated genes identified with RNA-seq in NPC1L1-sh C3H10 cells after induction with osteogenic medium for 3 days. F. RT-qPCR analysis of mRNA expression of cholesterol metabolism -related genes in osteoblast cells after induction with osteogenic medium for 3 days.

To further define the molecular mechanism underlying NPC1L1 modulate osteogenesis, we compared the whole transcriptome by RNA sequencing (RNA-seq) in control and NPC1L1-sh C3H10 cells after 0 or 3 days after osteogenic induction. Interestingly, when compared control C3H10 cells after 3days (con 3d) of induction with control C3H10 cells without induction (con 0d), gene ontology (GO) analyses revealed that genes in the cholesterol metabolic processes were significantly enriched in the up-regulation group (Figure 4D). On the contrary, when compared NPC1L1-sh C3H10 cells after 3days of induction (sh 3d) with NPC1L1-sh C3H10 cells without induction (sh 0d), gene ontology (GO) analyses revealed that genes in the cholesterol metabolic processes were significantly enriched in the down-regulation group (Figure 4E). In terms of cholesterol metabolism, to verify RNA-seq results, we performed RT-qPCR in the same experimental settings. The heatmap in Figure 4F exhibits the mRNA expression of cholesterol metabolic related genes, which all up-regulated in con 0d vs. con 3d group, and all down-regulated in sh 0d vs. sh 3d group, indicating NPC1L1 regulates cholesterol metabolism, but not cholesterol transportation, to inhibit osteogenic activity.

### 27-OHC generation mediates the osteogenesis inhibitive ability of NPC1L1

To determine the role of cholesterol metabolism in NPC1L1-induced osteoblast differentiation inhibition, we detected oxidation level of osteoblasts during osteogenesis via DCFH-DA, the fluorescent probe of ROS. Figure 5A shows that oxidation level of control C3H10 cells after 3 days of osteogenic induction was significantly higher than C3H10 cells without osteogenic induction, while NPC1L1-sh C3H10 cells had an obviously lower level of oxidation after 3 days of osteogenic induction. The statistical results of DCFH-DA positive cell percentage and mean fluorescence intensity (MFI) were consistent with the fluorescence images, indicating NPC1L1 may regulate generation of oxidation products of cholesterol during osteogenesis. To verify our hypothesis, metabolite products of cholesterol oxidation were tested using ELISA, and we found 27-hydroxycholesterol (27-OHC), one of the most abundant oxysterols, had a highly consistent concentration with oxidation level of osteoblasts in Figure 5A. And by exogenous addition of 27-OHC, could induce fluorescence production of DCFH-DA probes (Figure 5D and 5E). Next, we further validated the effect of 27-OHC on osteogenic differentiation. In osteoblasts with the addition of 27-OHC, we observed significant inhibition of osteogenic differentiation, accompanied by shallower AR-S and ALP staining (Figure 5F and 5G) and lower expression of osteogenic markers ALP, OPN, OSX, and OCN (Figure 5H), correspond to the results reported in the literature.

**Figure 5:**
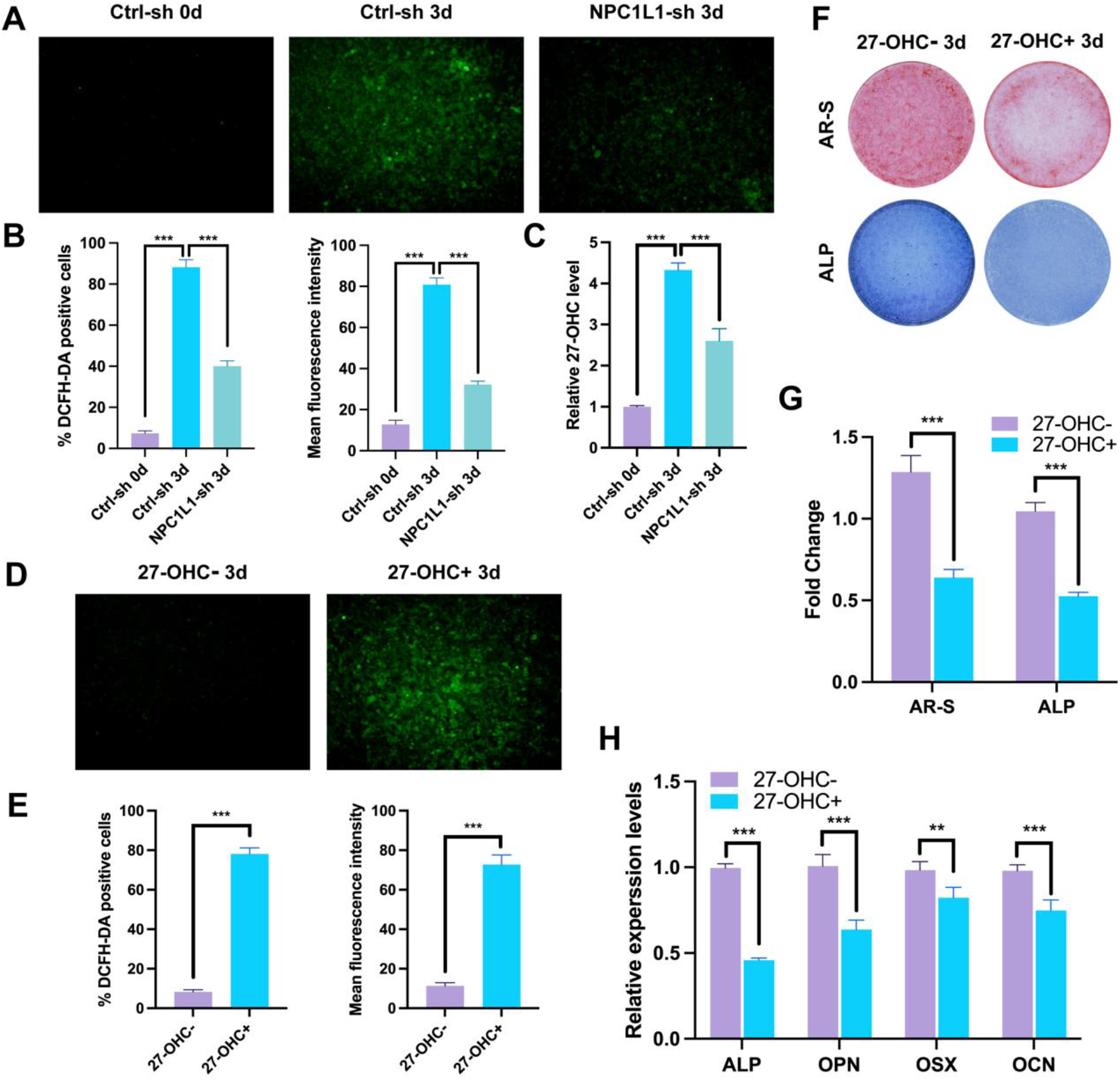
NPC1L1 induces 27-OHC accumulation during the process of osteoblast differentiation. A. Ctrl-sh and NPC1L1-sh C3H10 cells after induction with osteogenic medium for 3 days, as well as Ctrl-sh C3H10 cells without osteogenic medium, with ROS probe DCFH-DA treatment. B. Quantitative analysis of%DCFH-DA positive cells and mean fluorescence density in Figure 4A. C. 27-OHC level determined by ELISA. D. NPC1L1-sh cells, with or without 27-OHC treatment after induction with osteogenic medium for 3 days, with ROS probe DCFH-DA treatment. E. Quantitative analysis of%DCFH-DA positive cells and mean fluorescence density in Figure 4D. F. and G. AR-S and ALP staining of C3H10 cells with or without 27-OHC treatment after induction with osteogenic medium for 3 days. H. RT-qPCR analysis of osteogenic biomarker genes (ALP, OPN, OSX, and OCN) in C3H10 cells treated with or without 27-OHC. All bar graphs are presented as the mean ± SD. *P < 0.05; **P < 0.01; ***P < 0.001.

### NPC1L1 promotes Cyp27a1 expression through C/EBPα, thereby enhancing the generation of 27-OHC

To clarify how NPC1L1 controls cholesterol metabolism and 27-OHC generation, we found expression of Cyp27a1, a key metabolic enzyme that oxidizes cholesterol into 27-OHC, were significantly increased during normal osteoblast differentiation (Figure 4F and 6A) but down-regulated in sh 3d group when compared with con 3d in RNA-seq results (Figure 6B). RT-qPCR were performed and verified mRNA levels of Cyp27a1 were raised during osteogenesis, but lowered when NPC1L1 knockdown, suggested Cyp27a1 mediates the regulation of NPC1L1 on 27-OHC generation during osteoblast differentiation (Figure 6C and 6D). Therefore, we established NPC1L1 overexpression (NPC1L1-OE) osteoblasts and conducted osteogenesis induction and staining experiment. As we expected, NPC1L1 overexpression inhibited osteogenesis, but Dafadine-A, the specific inhibitor of Cyp27a1, obviously reversed the inhibition effect of NPC1L1 on osteoblast differentiation (Figure 6E, 6F, and S3). Similarly, mRNA expression of osteogenic biomarkers reduced significantly in NPC1L1-OE group but had a relatively close to normal levels when treated with Dafadine-A, indicating NPC1L1 regulated Cyp27a1 mRNA expression to enhance 27-OHC production (Figure 6). According to the promoter region sequence of Cyp27a1, we predicted potential upstream transcription factors via JASPER and UCSC database. Combining the RNA-seq results in this subject, we speculate that C/EBPα most likely to mediate the regulatory effect of NPC1L1 on Cyp27a1 expression. Then, CHIP-qPCR were performed using control and NPC1L1-OE osteoblasts and verified C/EBPα binds to the promoter region of Cyp27a1 and promotes its expression. We further validate the above results at the protein level, representative western blot analysis showed NPC1L1 increases C/EBPα level to up-regulate Cyp27a1 expression, further raise the concentration of 27-OHC to inhibit osteogenic differentiation. Finally, rescue experiment was performed using MTL-C/EBPα to reverse the down-regulated C/EBPα in NPC1L1-sh osteoblasts, it can be seen that the previously decreased Cyp27a1 increases accordingly. Taken together, transcription factor C/EBPα mediates the promoting effect of NPC1L1 on Cyp27a1 expression in osteoblasts, further elevated 27-OHC level to inhibit osteogenesis.

**Figure 6:**
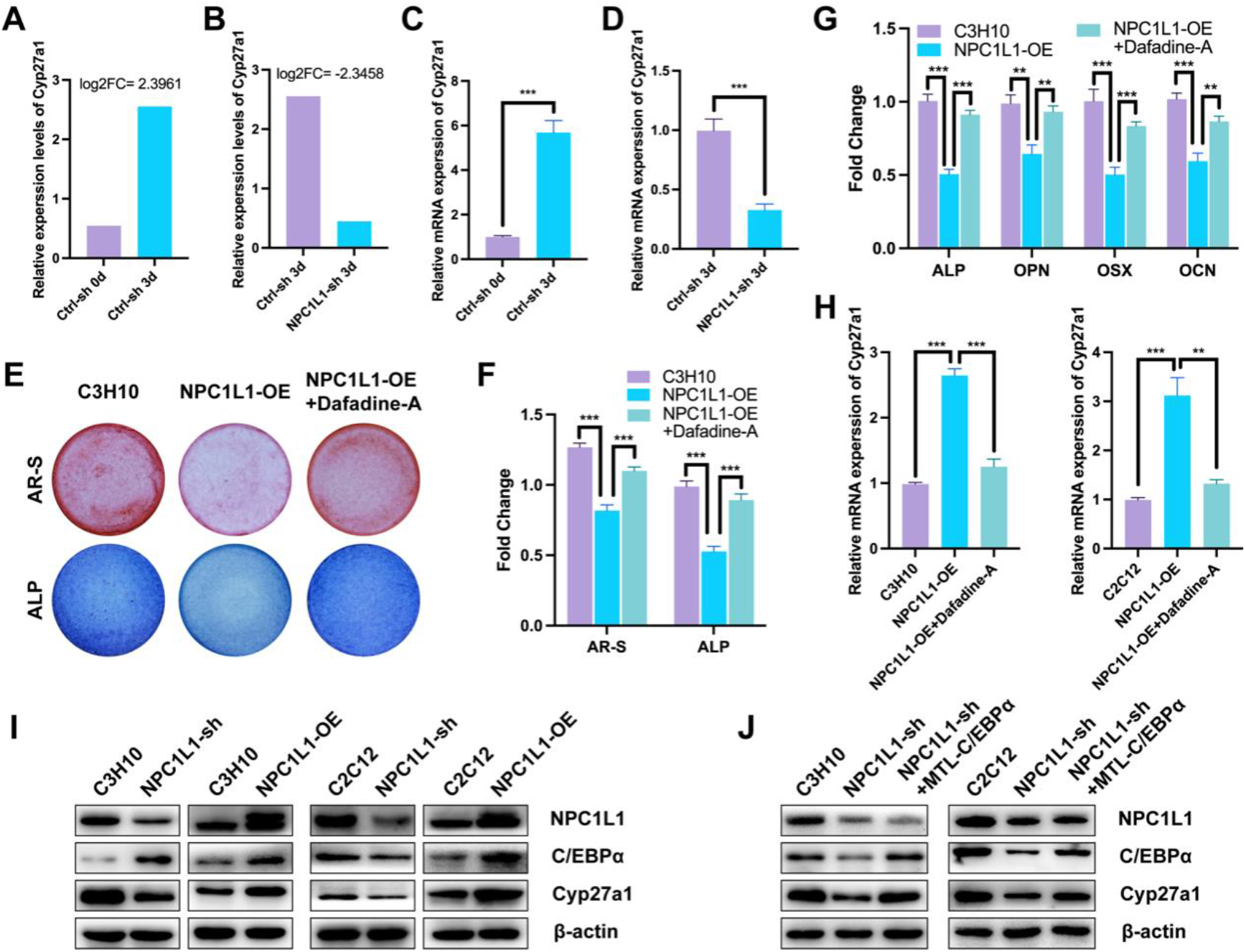
C/EBP*α*/Cyp27a1/27-OHC axis mediates NPC1L1-induced osteogenesis inhibition. A. and B. Relative expression level of expression in RNA-seq results of Ctrl-sh or NPC1L1-sh cells after induction with osteogenic medium for 3 days. C. and D. RT-qPCR analysis of mRNA expression of Cyp27a1 expression in RNA-seq results of Ctrl-sh or NPC1L1-sh cells after induction with osteogenic medium for 3 days. E. and F. AR-S (upper panel) and ALP (lower panel) staining of control, NPC1L1 overexpression (NPC1L1-OE), and Dafadine-A treated NPC1L1-OE C3H10 cells after induction with osteogenic medium for 3 days. G. RT-qPCR analysis of osteogenic biomarker genes (ALP, OPN, OSX, and OCN) in control, NPC1L1 overexpression (NPC1L1-OE), and Dafadine-A treated NPC1L1-OE C3H10 cells after induction with osteogenic medium for 3 days. H. Relative mRNA expression of Cyp27a1 in control, NPC1L1 overexpression (NPC1L1-OE), and Dafadine-A treated NPC1L1-OE C3H10 cells. I. Representative western blot analysis of NPC1L1, C/EBPα, and Cyp27a1 in NPC1L1-sh or NPC1L1-OE osteoblast cells. J. Representative western blot analysis of NPC1L1, C/EBP*α*, and Cyp27a1 in control, NPC1L1-sh, and MTL-C/EBPα treated NPC1L1-sh osteoblast cells. All bar graphs are presented as the mean ± SD. *P < 0.05; **P < 0.01; ***P < 0.001.

## Discussion

Cholesterol is an important component of the membrane structure of cells, and also an important raw material for synthesizing steroids, oxysterols, and bile acids, which involves in various physiological activities of the human body and cells. Recently, an increasing number of studies have focused on the important regulatory role of cholesterol metabolism in bone formation and osteoblast differentiation, but the exact mechanism still needs to be clarified[25]. For example, loss of Sc5d (Sc5d-/-), an enzyme that catalyzes of lathosterol into 7-dehydrocholesterol, would results in cleft palate, micrognathia, and abnormal limb formation. Specifically, Sc5d-/-mice exhibited defective osteoblast differentiation and mandibular hypoplasia[26]. Besides, decreased expression of Dhcr7, another cholesterol biosynthesis regulating enzyme, causing cholesterol metabolism abnormalities and osteoblast differentiation dysfunction during the process of osteogenesis[27]. The above researches demonstrate that cholesterol metabolism and osteogenic differentiation compensation may be involved, with the former having an important regulatory relationship with the latter. However, the exact role of cholesterol metabolism in osteoblast differentiation remains unknown. In our previous study, osteogenesis was found regulated by circadian rhythm in the context of osteoporosis, and then RNA-seq was conducted to elucidate the potential molecular mechanisms[28]. As is shown in Figure 1A, lipid and cholesterol metabolism were changed remarkably, and the key cholesterol transporter NPC1L1 was one of the molecules with the most significant changes (Figure 1B and 1C), which verified the role of cholesterol metabolism in osteoblast differentiation and inspired us to clarify the mechanism of how NPC1L1 regulates cholesterol to control osteogenesis.

NPC1L1 was reported that plays a vital role in intestinal cholesterol absorption, dietary cholesterol absorption and biliary cholesterol resorption via mediating cholesterol absorption into cells. However, few studies have focused on the function of NPC1L1 protein in regulating cholesterol metabolism. In this study, we firstly explored whether NPC1L1 regulates osteogenesis through mediating cholesterol entry into cells as a transporter. Interestingly, Eze didn’t affecting osteoblast differentiation, as it blocks cholesterol transport function of NPC1L1. Through comparing NPC1L1 knockdown cells with control cells via another round of RNA-seq, it is worth noting that all enriched cholesterol metabolism-related pathways were upregulated when comparing control osteoblasts after 3 days of osteogenic induction with 0 day of induction (Figure 4D), but all enriched cholesterol metabolism-related pathways were downregulated when comparing NPC1L1-sh osteoblasts after 3 days of osteogenic induction with 0 day of induction (Figure 4E). On the one hand, consistent with literatures, our data indicates that cholesterol metabolism is closely related to osteogenic differentiation, and shows an activation trend in the early stages of osteogenesis (Figure 4D). On the other hand, cholesterol metabolism was significantly inhibited during osteogenesis process when NPC1L1 downregulated, suggesting NPC1L1 regulates osteogenic differentiation by regulating cholesterol metabolism rather than transportation. At present, there is no literature reporting the osteogenic regulating ability of by NPC1L1, and research related to NPC1L1 mainly focuses on cholesterol metabolism in the intestine and liver. Therefore, this article will mainly study the role and mechanism of NPC1L1 in regulating osteoblast differentiation, expected to elucidate the mechanism of NPC1L1 mediated cholesterol metabolism in the occurrence and development of osteoporosis.

Oxysterols are derived from cholesterol, exists in different parts of the human body, and has rich biological functions. For example, circulating 7α-hydroxycholesterol (7α-OHC) is catalyzed by Cyp7a1 and is a starting intermediate in the biosynthesis of bile acids; 24(S)-Hydroxycholesterol (24-OHC) is the main metabolite of brain cholesterol and plays a major role in maintaining the homeostasis of cholesterol in the brain; 20(S)-hydroxycholesterol (20S-OHC) is an activator of the hedgehog (Hh) signaling to induce osteogenic differentiation of BMSCs while also inhibiting their adipogenic differentiation; and 27-hydroxycholesterol (27-OHC) is catalyzed by the mitochondrial enzyme sterol 27-hydroxylase (Cyp27a1), which is a selective estrogen receptor modulator and an agonist of the liver X receptor widely distributed in tissues. 27-OHC is a negative regulator of bone homeostasis, which impairs bone formation and promotes bone turnover through the activation of LXR and upregulation of the osteoblast TNF-α and RANKL expressions[29, 30]. Moreover, 27-OHC was reported that it can increase cellular reactive oxygen species (ROS) levels and inhibit cellular viability. ROS plays dual role in osteoblast differentiation, as it acts as signaling molecules participates in cellular differentiation and proliferation at the physiological levels, but also leads to damage to lipids, proteins, and DNA, which result in apoptosis and autophagy at the cumulative levels. In this study, we detected the oxidative stress levels during osteogenic differentiation via DCFH-DA, and verified the ROS accumulation. Subsequently, cholesterol metabolites were tested and found 27-OHC significantly upregulated during osteogenesis, but had an unconspicuous elevation in NPC1L1-sh osteoblasts (Figure 5A-C). On the contrary, the artificial addition of 27-OHC can also increase the ROS level of NPC1L1-sh cells during osteogenic differentiation, and inhibiting differentiation ultimately. Therefore, we suppose that NPC1L1 promotes cholesterol metabolism to generate 27-OHC, which elevates ROS levels and inhibits osteogenic differentiation. Conversely, inhibiting NPC1L1 reduces 27-OHC levels and promotes differentiation.

As is mentioned above, 27-OHC is generated from cholesterol by the sterol hydroxylase Cyp27a1, which is a member of the cytochrome p450 superfamily. Cyp27a1 may have a regulatory effect on bone homeostasis, as female Cyp27a1-/-mice exhibited more trabeculae than Cyp27a1+/+ mice, but these trabeculae are thinner[31, 32]. Besides, Fang et al. reported that Cyp27a1 deficiency enhanced osteoclast differentiation and bone loss[33]. The mechanism involved and the regulatory effect of Cyp27a1 on osteoblasts are not yet clear, and the main consideration is to regulate the generation of 27-OHC. Through analyzing the transcriptomic data of C3H10 cells before and after osteogenic induction, expression level of Cyp27a1 obviously upregulated, consistent with the expression trend of 27-OHC (Figure 4F). Although there are no significant changes of Cyp27a1 expression between NPC1L1-sh osteoblasts before and after osteogenic induction, we speculate that Cyp27a1 is in a sustained low expression state after knocking down NPC1L1, while the Cyp27a1 expression level of NPC1L1-sh cells was significantly lower than that of control cells after osteogenic induction. The enhanced osteogenic staining and expression of osteogenic biomarkers significantly of NPC1L1-OE cells after treating with Dafadine-A, the specific inhibitor for Cyp27a1, confirming the hypothesis that NPC1L1 promotes 27-OHC levels to inhibit osteogenic differentiation by activating Cyp27a1. C/EBP*α* is one of the master regulators of adipogenesis and controls the balance between osteogenic and adipogenic differentiation during OP. To further clarify the mechanism by which NPC1L1 promotes Cyp27a1, the data shows that both the mRNA and protein levels of Cyp27a1 have changed, suggesting that NPC1L1 promotes the transcription of Cyp27a1. Based on the promoter sequence of Cyp27a1, we used the JASPAR database to predict and analyze that CEBPA may be the upstream transcription factor of Cyp27a1, so we designed a series of experiments to validate it. Our results exhibit that C/EBPα directly binds to the Cyp27a1 gene and enhanced its expression, while NPC1L1 significantly promotes the expression of C/EBPα. Moreover, activating the expression of C/EBPα through MTL-C/EBPα restore the expression of previously downregulated Cyp27a1 in NPC1L1-sh cells.

## Conclusion

In conclusion, our study identifies the previously unknown osteogenic inhibitory role of NPC1L1 in regulating 27-OHC generation through C/EBPα/Cyp27a1 axis during OP. Therefore, NPC1L1, as well as C/EBPα/Cyp27a1/27-OHC axis, could be an important therapeutic target for the treatment of OP. Our results suggesting a cholesterol regulatory function for NPC1L1 besides its activity as a (re-)uptake transporter.

## Acknowledgement

This work was sponsored by Natural Science Foundation of Shanghai (22ZR1448900); National Natural Science Foundation of China (82303139); the China National Postdoctoral Program for Innovative Talents (BX20230084); China Postdoctoral Science Foundation (2023M740687); Shanghai Pujiang Program (No. 22PJD010); Natural Science Foundation of Minhang District, Shanghai (2021MHZ081, 2020MHZ028); the Key Department of Minhang District, Shanghai (2020MWTZB03); the Key Department of the Fifth People’s Hospital of Shanghai (2020WYZDZK03); the Fifth People’s Hospital of Shanghai, Fudan University (N123E5).

## Supplementary File

